# Influence of viral transport media and freeze-thaw cycling on the sensitivity of qRT-PCR detection of SARS-CoV-2 nucleic acids

**DOI:** 10.1101/2021.06.18.448982

**Authors:** Cian Holohan, Sophia Hanarahan, Nathan Feely, Peng Li, John O’Connell, Catherine Moss, Michael Carr, Oya Tagit, Gil U Lee

## Abstract

The events of the last year have highlighted the complexity of implementing large-scale molecular diagnostic testing for novel pathogens. The purpose of this study was to determine the chemical influences of sample collection media and storage on the stability and detection of viral nucleic acids by qRT-PCR. We studied the mechanism(s) through which viral transport media (VTM) and number of freeze-thaw cycles influenced the analytical sensitivity of qRT-PCR detection of SARS-CoV-2. Our goal is to reinforce testing capabilities and identify weaknesses that could arise in resource-limited environments that do not have well-controlled cold chains. The sensitivity of qRT-PCR analysis was studied in four VTM for synthetic single-stranded RNA (ssRNA) and double-stranded DNA (dsDNA) simulants of the SARS-CoV-2 genome. The sensitivity and reproducibility of qRT-PCR for the synthetic ssRNA and dsDNA were found to be highly sensitive to VTM with the best results observed for ssRNA in HBSS and PBS-G. Surprisingly, the presence of epithelial cellular material with the ssRNA increased the sensitivity of the qRT-PCR assay. Repeated freeze-thaw cycling decreased the sensitivity of the qRT-PCR with two noted exceptions. The choice of VTM is critically important to defining the sensitivity of COVID-19 molecular diagnostics assays and this study suggests they can impact upon the stability of the SARS-CoV-2 viral genome. This becomes increasingly important if the virus structure is destabilised before analysis, which can occur due to poor storage conditions. This study suggests that COVID-19 testing performed with glycerol-containing PBS will produce a high level of stability and sensitivity. These results are in agreement with clinical studies reported for patient-derived samples.

## Introduction

The ongoing coronavirus disease 2019 (COVID-19) pandemic caused by a novel strain of severe acute respiratory syndrome coronavirus, SARS-CoV-2, has claimed almost 4 million lives worldwide as of June 2021.^1^ Efficient management of the pandemic requires rapid and accurate identification and isolation of infected symptomatic and asymptomatic individuals at the early stages of infection.

The current gold standard for COVID-19 diagnosis is quantitative real-time reverse transcription polymerase chain reaction (qRT-PCR)-based detection of viral RNA obtained from patient samples (Figure 1A).^2^ The qRT-PCR assay is a highly sensitive method with a limit of detection of single RNA copy.^3^ However, the sensitivity and accuracy of qRT-PCR can be hindered by low viral loads particularly when screening patients in the pre-symptomatic phase^4^, and by contamination.^5,6^ In this regard, RNA extraction kits utilizing lysis buffers and silica-coated magnetic beads can significantly improve qRT-PCR detection sensitivity by facilitating an efficient release, capture, and isolation of viral RNA from the infected cells, thereby providing purified and concentrated RNA samples suitable for qRT-PCR analysis (Figure 1B).^7-9^ Maintaining the stability of patient-derived samples during their collection, storage and transport to diagnostic facilities is equally important for sensitive and accurate detection of SARS-CoV-2 as improper storage and transport conditions can cause the degradation of labile viral RNA and subsequently lead to false negatives with serious implications for the tracking and tracing of virus outbreaks.^10,11^

**Figure 1.**
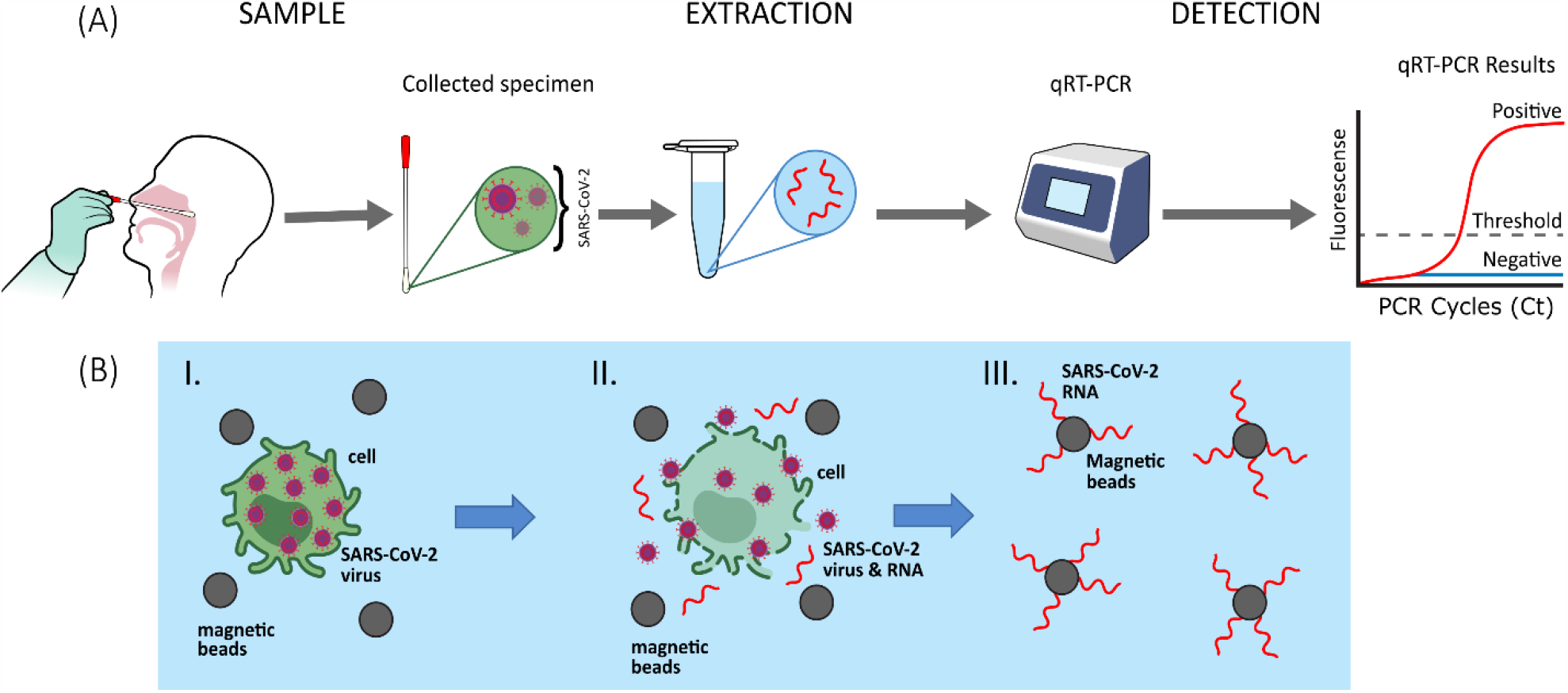
**(A)** Schematic illustration of the molecular diagnostics workflow for the detection of SARS-CoV-2 RNA from patient samples using qRT-PCR. The nasopharyngeal swab obtained from the patient is stored in viral transport media (VTM) while in transit to a testing facility. The sample is then processed using an RNA extraction procedure to isolate viral RNA prior to qRT-PCR amplification. **(B)** An illustration of the RNA extraction process. (I) Cells containing SARS-CoV-2 viral particles are disrupted in lysis buffer containing silica-coated magnetic beads, denaturants and detergents. (II) SARS-CoV-2 RNA released from the lysate is captured by silica-coated magnetic beads by adsorption. (III) Magnetic separation of adsorbed RNA allows for the elimination of cellular debris and contaminants before the RNA is finally eluted into RNase-free water.

While SARS-CoV-2 has been detected in several (liquid) biopsies such as saliva, blood, tears, urine and feces in a range of viral loads and with varying degrees of stability, ^12-14^ sample collection from the upper and lower respiratory tract using nasopharyngeal swabs is the standard practice for diagnostic testing for respiratory prathogens.^15,16^ Immediately after collection, the nasopharyngeal swabs are deposited into sterile tubes containing viral transport media (VTM) to preserve the sample until diagnostic tests are performed. VTM contain various proteins, amino acids and antimicrobial agents suspended in a buffered salt solution. These can be purchased commercially, e.g., universal transport media, UTM^®^, or prepared using recipes provided by the US Centre for Disease Control and Prevention (CDC) ^17^ and the World Health Organization (WHO).^18-20^ The importance of VTM selection for accurate qRT-PCR-based molecular diagnostics has been highlighted in a recent study by Kirkland and Frost (2020). It was shown that the composition of commercially available VTM may not always be suitable for the intended purpose, such as nucleic acid detection, which is not always obvious to clinicians and medical scientists.^21^

In addition to VTM composition, the storage and transport temperature of patient-derived samples can significantly affect virus stability, and thus the accuracy of diagnostic testing. SARS-CoV-2 is highly stable at temperatures up to 4°C, but is sensitive to elevated temperatures.^22^ The lipid bilayer in the enveloped structure of SARS-CoV-2 makes it more susceptible to heat inactivation compared to non-enveloped viruses.^23^ Therefore, it is recommended that samples are kept at 2-8°C and transported to a diagnostic lab within 72 h after collection. If this is not possible, samples should be stored at -70°C for periods longer than 72 hr.^15^ Failure to maintain appropriate cold chains can result in repeated freeze-thawing of samples, which could degrade the viral RNA and potentially cause false-negative reporting.^11,24,25^

Although separate studies have examined the influence of VTM selection^21,26^ and repeated freeze-thaw cycles^27, 25^ on the stability of SARS-CoV-2 samples, the combined effects of different VTM compositions and the repeated freeze-thaw cycles on sample stability have not been shown. The present study aims to determine how repeated freeze-thawing affects the stability of samples with varying quantities of nucleic acids in different VTM, and to suggest improvements in sample collection, storage, and transport conditions to improve SARS-CoV-2 molecular diagnostics. To mimic the viral genome extracted from patient samples, we employed synthetic linear SARS-CoV-2 RNA fragments made up of six non-overlapping 5 kb single-stranded RNA (ssRNA) units that cover 99.9% of the Wuhan-Hu-1 reference genome (GeneBank ID: MN908947.3). In parallel, a double-stranded DNA (dsDNA) plasmid (ca. 4 kb in size) harbouring the SARS-CoV-2 *nucleocapsid* (*N*) gene was tested as a more stable control less susceptible to nuclease degradation. Nucleic acid extraction and subsequent qRT-PCR allowed us to investigate how different VTM and the freeze-thaw process differentially affect the two synthetic SARS-CoV-2 products and, also, in the presence of epithelial cells to achieve a more accurate mimicry of patient-derived samples.

## Experimental

### Materials

All reagents and kits were commercially available and used as received without further alterations, unless specified otherwise. The viral RNA extraction kit containing silica-coated high magnetisation beads and magnetic separator were developed as reported previously.^29,30^ DNase-and RNase-free hydrophobic filtered pipette tips, 96-well plates, sealing covers and microtubes were used throughout. Environmental and sample RNase activity levels were monitored throughout the study using the RNaseAlert^®^QC system V2 (ThermoFisher, US). Copan universal transport medium (UTM^®^) 330 C was obtained from Medical Supply Company Ltd., Ireland. Hanks’ balanced salt solution (HBSS), glycerol, heat inactivated fetal bovine serum (FBS), gentamicin, and amphotericin B were purchased from ThermoFisher, US. Penicillin G sodium salt S, streptomycin, polymyxin B, nystatin, moxifloxacin and sulfamethoxazole were from Sigma, UK. All sample preparation and handling were carried out in an aseptic manner in a sterile, RNase-free BSL-2 facility.

### Preparation of viral transport media

HBSS VTM was prepared by adding gentamicin (50 mg) and amphotericin B (0.25 mg) to sterile HBSS (500 mL). The solution was thoroughly mixed and stored at 2-8 °C. CDC VTM was prepared as per CDC guidelines.^17^ In brief, gentamicin (50 mg), amphotericin B (0.25 mg) and FBS (10 mL) were added to sterile HBSS (500 mL). The solution was thoroughly mixed and stored at 2-8 °C. WHO VTM was prepared as per WHO guidelines.^18^ Briefly, penicillin G sodium salt (1,000,000 I.U.), streptomycin (100 mg), polymyxin B (1,000,000 I.U.), gentamicin (125 mg), nystatin (250,000 I.U.), moxifloxacin (30 mg) and sulfamethoxazole (100 mg) were added to a sterile solution of 1:1 PBS: glycerol (500 mL), filtered using 0.45 µm pore-sized membranes, thoroughly mixed and stored at -20 °C.

### Monitoring RNAse activity

A fluorometric RNaseAlert^®^ kit was employed to measure RNase activity in all prepared buffers and VTM. The duplicate measurements were performed for each sample in 96-well optical plates as per the operating instructions. The samples were monitored with a ClarioStar Plus fluorescence plate reader over a 35-45 min incubation at 37°C with orbital mixing at 500 RPM (excitation/emission 490/520 nm). A mean fluorescence intensity two-fold higher than that of the negative control was defined as positive for RNase activity.^31^

### Preparation of SARS-CoV-2 nasopharyngeal swab mimics

Commercially available synthetic nucleic acid fragments of SARS-CoV-2 that are routinely used as positive controls in qRT-PCR molecular diagnostics for SARS-CoV-2 were used as viral mimics in this study. These consisted of the Twist Biosciences ssRNA control (GenBank ID: MN908947.3, GISAID: WUHAN-HU-1) and a dsDNA plasmid control obtained from Integrated DNA Technologies (IDT). Samples were prepared in 7,000, 70,000 and 700,000 viral copy number (VCN)/mL viral loads by spiking 1 mL of VTM with corresponding number of copies of genetic material. To further mimic a standard nasopharyngeal swab that also contains cells taken from the nasopharynx, A375 epithelial human melanoma cells (ATCC^®^ CRL-1619 ™) counted by hemocytometry were mixed with SARS-CoV-2 ssRNA in a 1:10 cell to RNA ratio (70,000, 7,000 and 700 cells/mL).

### Freeze-thaw treatment of SARS-CoV-2 nasopharyngeal swab mimics

The swab mimics were prepared as 4.5 mL stocks in RNase-free cryotubes as described above. Each stock was frozen in a -80 °C freezer for 4 h. To retrieve aliquots for qRT-PCR analysis, the stocks were thawed at room temperature for 1 h and the remaining stock was then returned to the -80 °C freezer to begin a new freeze-thaw cycle. This process was repeated up to 10 freeze-thaw cycles.

### Preparation of lysis and wash buffers

Lysis and wash buffers were prepared in-house and tested for RNase contamination prior to use in RNA extraction experiments. A basal lysis buffer containing 50 mM Tris, 6 M guanidine thiocyanate, 25 mM EDTA was prepared, and pH was adjusted to 6.5 using HCl. To prepare the lysis buffer, the basal lysis buffer was supplemented with 3% (w/v) Triton X-100. Wash buffer 1 contained 50 % (v/v) ethanol and basal lysis buffer. Wash buffer 2 contained 80 % (v/v) ethanol and nuclease-free water.

### Sample lysis and RNA purification

RNA extraction was conducted with silica-coated magnetic beads in a 96-well plate using the M96D-400 magnetic separator (Magnostics Ltd) under aseptic, RNase-free conditions. 1,4-dithiotheritol (DTT, 2% w/v) and polyadenylic acid carrier RNA (poly (A), 2 mg/mL) were added to the lysis buffer. 250 µL of sample containing 1,750, 17,500, or 175,000 viral copies were treated with lysis buffer at room temperature for 10 min, which was followed by the addition of the silica-coated magnetic beads and 250 µL of isopropanol. The mixture was shaken vigorously for 10 min and washed sequentially with wash buffers 1 and 2 to remove impurities. The purified RNA was eluted from the magnetic beads into 50 µL of nuclease free water and stored at -80 °C for up to 5 days prior to qRT-PCR analysis. A375 cells were run in parallel with all extractions and qRT-PCR analysis of the purified nucleic acid material for the human *glyceraldehyde 3-phosphate dehydrogenase* (*GAPDH*) endogenous control gene was treated as a positive extraction control. All extractions were performed in triplicate.

### Quantitative reverse transcription - polymerase chain reaction (qRT-PCR) analysis

A master-mix was created using the SuperScript™ III Platinum™ One-Step qRT-PCR kit (Invitrogen, US) in conjunction with N1 primers/probes from the IDT 2019-nCov CDC EUA Kit (IDT, US) with the following composition: SuperScript™ III RT/Platinum™ Taq mix (16 µL), 2X reaction mix (400 µL), ROX™ reference dye (1.6 µL), IDT N1 primers/probes (60 µL; final concentration of 800 nM primers and 100 nM of FAM-labelled probes) and nuclease free water (122.4 µL). 15 µL aliquots of the master mix was used per 5 µL of the extracted samples to detect both the ssRNA and dsDNA plasmid using qRT-PCR. Therefore 175, 1,750 and 17,500 viral copies were present in the qRT-PCR reaction from the initial 7,000, 70,000 and 700,000 VCN/mL stocks, respectively. A second master mix for the qRT-PCR analysis of the *GAPDH* gene for both the A375 extraction controls run in tandem with cell-free RNA spiked VTM was created as described above using *GAPDH*-specific primers and probe (40 µL; final concentration of 10 nM primers and 17 nM probes of VIC-labelled probe) in place of N1 primers/probe and nuclease free water (142.4 µL).

The samples were kept on ice immediately prior to qRT-PCR analysis using a QuantStudio™ 7 Flex Real-Time PCR (Applied Biosystems^®^, US). The run parameters followed a standard PCR cycle beginning with reverse transcription at 50 °C for 15 min, denaturation at 95 °C for 2 min followed by 50 amplification cycles consisting of a denaturation step at 95 °C for 15 s and annealing and extension at 60 °C for 1 min with fluorescence acquisition in the annealing/extension phase. QuantStudio Software v1.3 was used to analyze the data. The cycle threshold (Ct) was set at 0.05 and baseline set to automatic. Two-tailed t-tests were performed using GraphPad software in order to make comparisons between samples. These comparisons were regarded statistically significant for p < 0.01.

## Results and Discussion

### Influence of VTM on SARS-CoV-2 detection sensitivity using qRT-PCR

The analytical sensitivity of qRT-PCR detection was studied using a synthetic single-stranded RNA (ssRNA). To determine the influence of RNase activity on the detection sensitivity, a synthetic double-stranded DNA plasmid was tested in parallel. The ssRNA construct consisted of six non-overlapping 5 kb nucleotide sequences that covered 99.9% of the SARS-CoV-2 Wuhan-Hu-1 reference genome in a state that is not protected by the viral capsid and thus susceptible to both chemical and physical degradation. The DNA construct was a ∼4 kb plasmid that includes the SARS-CoV-2 *N* gene. The impact of the nasal membrane cellular materials on the qRT-PCR ssRNA sensitivity was studied by addition of A375 epithelial cells to ssRNA in a 1:10 ratio.

qRT-PCR relies on fluorescent reporters to monitor the amplification of target nucleotide sequences for each thermal cycle. An amplification plot with the relative fluorescence intensity (log Δ*Rn*) is plotted as a function of thermal cycle number to detect the target sequences. A representative amplification plot is shown in Figure 2A for three samples: a positive *GAPDH* control for A375 human epithelial cells (green), a sample with positive amplification comprising 17,500 copies per qRT-PCR reaction after extraction from PBS-G VTM with cells, (orange), and an undetected sample below the assay limit of detection comprising 1,750 copies per qRT-PCR reaction after extraction from CDC VTM with cells (red). The *Rn* values were obtained by dividing the fluorescence of the reporter dyes (either N1 or *GAPDH* assays) by the fluorescence of a passive reference dye (ROX™). The *ΔRn* values were determined by subtracting the *Rn* value of the baseline signal from that of the experimental reaction. Wölfel et al reported an average viral load per swab in 3 mL of VTM for 19 patients to be 676,000 viral copies. In this study we use a VTM volume per sample condition of 1 mL while using 700,000 VCN as our highest copy number. ^32^

**Figure 2.**
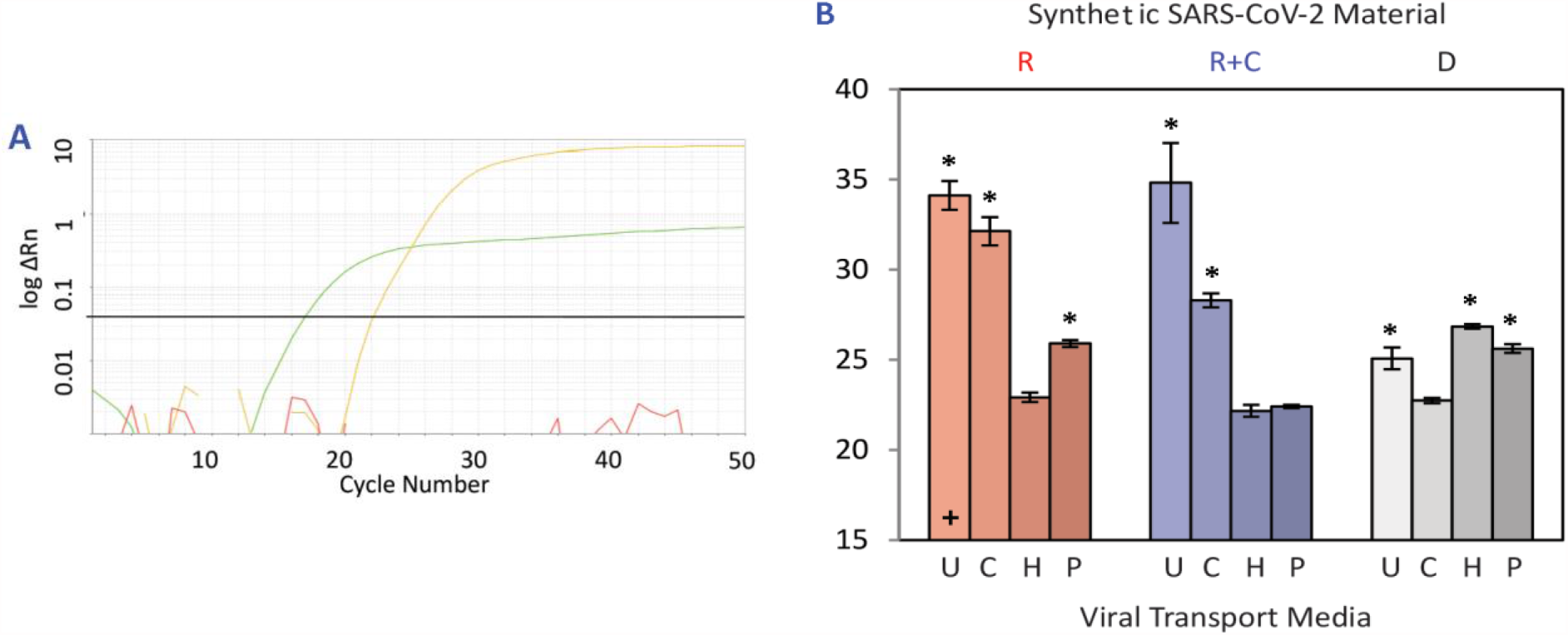
Impact of viral transport media on detection of synthetic SARS-CoV-2 nucleic acids by qRT-PCR. **(A)** Amplification plot of the qRT-PCR results, depicting the log*ΔRn* plotted against the cycle number. Positive extraction control curve from human epithelial A375 cells (green) amplified with *GAPDH* primers/probe, a positive signal derived from 17,500 copies per qRT-PCR reaction of ssRNA+cells extracted from PBS-G VTM (orange), and an undetected reaction containing 1,750 copies per qRT-PCR reaction of ssRNA+cells extracted from CDC VTM (red) both amplified by N1 primers/probe are shown. The threshold set at 0.05 is indicated by the horizontal line (black). **(B)** Results for ssRNA (R), ssRNA+cells (R+C) and dsDNA (D) sample types at 17,500 copies per qRT-PCR reaction extracted from UTM(U), CDC (C), HBSS (H) and PBS-G (P) VTM. Measurements were performed in triplicate. The number of failed amplifications per triplicate measurement is denoted by a cross (+) per failed run, i.e., one cross indicates one failed run. Two-tailed t-tests were used to compare the VTM that produced the best result for each sample type (H for sample R, H for sample R+C, and C for sample D) to the other VTM results of that sample type. Samples that produced a statistically significant difference (p < 0.01) are denoted by the presence of an asterisk (^*^).

Figure 2B presents the qRT-PCR results for the three sample types, i.e., ssRNA (R), ssRNA+cells (R+C), and dsDNA (D) extracted from UTM, CDC, HBSS and PBS-G VTM at 17,500 copies per qRT-PCR reaction. After multiple thermal cycles the target sequence is amplified to a point whereby the fluorescence signal emitted crosses a threshold for detection. The cycle number at which this crossing the threshold occurs is defined as the *Ct* value. In this study the threshold was set at a Δ*Rn* value of 0.05. A lower *Ct* value indicates the presence of a larger quantity of initial target sequence in the sample as fewer thermal cycles are required to cross the threshold and vice-versa. A trend that can be seen for the ssRNA results (R) is that the *Ct* values are lower in the HBSS (22.92, σ = 0.25) and PBS-G (25.90, σ = 0.19) VTM compared to the UTM (34.13, σ = 0.8) and CDC (32.13, σ = 0.79) VTM (Figure 2B). A similar trend is observed in the presence of A375 epithelial cells (R+C), such that the lowest and highest *Ct* values are obtained for HBSS (22.16, σ = 0.33) and UTM (34.82, σ = 2.22), respectively. Overall, a subtle decrease in *Ct* values (increased assay sensitivity) is noted for all VTM when epithelial cells are present (R+C), with the exception of UTM. Even though sample type R with U VTM produced a lower *Ct* than sample type R+C U, one R U triplicate failed to amplify while there were no failed amplifications for R+C U samples.

The influence of the VTM on dsDNA qRT-PCR results (D) was less apparent and does not follow the same trend seen in the R and R+C data, whereby UTM and CDC VTM produced higher *Ct* values than HBSS and PBS-G. In contrast, sample type D C (22.74, σ=0.14) and D U (25.08, σ=0.6) produced lower *Ct* values than D P (25.62, σ=0.24) and D H (26.85, σ=0.12). However, all D qRT-PCR *Ct*s were significantly higher (p < 0.01) compared to the best results obtained in CDC VTM. These results highlight the important role that the VTM selection has when using different synthetic SARS-CoV-2 nucleic acids and how it can affect the qRT-PCR assay sensitivity.

An RNase activity assay was performed to characterise presence of nucleases in the VTM, and both the CDC and UTM VTM tested positive for RNase activity (Figure S1, Supplementary Information). Each component of the CDC VTM was tested to identify the source of the RNase activity and FBS was the only component which tested positive for the presence of RNases. The FBS sample showed a high fluorescence intensity of ∼ 10,000 A.U. from the first time point (90 s) and therefore was considered to have a high level of RNase activity (Figure S2, Supplementary Information). The positive control supplied with the kit was RNase A, which had a fluorescence intensity of ∼11,500 A.U. by the end of the incubation period. This led us to conclude that FBS was the source of RNase activity in this VTM. The Copan UTM also tested positive for RNase activity with an average fluorescence intensity of 10,000 A.U. at the end of the incubation period, with the positive RNase control averaging 11,500 A.U. by the end of the incubation period (Figure S1, Supplementary Information). Both HBSS and PBS-G VTM as well as all the reagents used in the extraction were found to be RNase free (Figure S1 and S3, Supplementary Information).

The RNase activity assays results provide insight into why samples containing ssRNA and ssRNA+cells produced poor results in CDC and UTM when compared to their HBSS and PBS-G counterparts. HBSS VTM preparation is identical to CDC VTM preparation, but excludes the addition of the RNase-containing FBS. In addition, qRT-PCR detection of A375 epithelial cells in ssRNA+cells samples after extraction from VTM helped to assess whether the presence of RNases in the VTM similarly affected the cells in these samples. Interestingly, cells at 1,750 copies per qRT-PCR reaction extracted from CDC (22.17, σ=0.26) and UTM (23.96, σ=0.18) produced higher *Ct*s than cells extracted from HBSS (18.56, σ=0.18) and PBS-G (18.85, σ = 0.13). Taking these findings together, it is evident that the VTM which tested positive for RNase activity had a largely negative effect on qRT-PCR detection sensitivity for all samples containing ssRNA. These results are in line with those of Kirkland and Frost ^21^, who also showed the adverse effects of the presence of FBS in VTM and how the use of such a VTM may impact results in molecular diagnostic and research applications. In contrast to these findings, dsDNA samples in CDC and UTM produced better results compared to dsDNA in HBSS and PBS-G, suggesting that the presence of FBS positively impacted on the dsDNA plasmid. This may be attributable to dsDNA being inherently more stable than ssRNA as the latter is more labile and has greater exposure to RNases in the environment. ^33^

Surprisingly, lower *Ct* values were obtained for all VTM tested in the presence of epithelial cells, except UTM (R+C, Figure 2b), implicating a better stabilisation and a more efficient extraction of ssRNA when performed with cellular nucleic acids. Indeed, the RNase activity assay performed on the RNase-positive VTM with and without cells showed a reduced RNase activity when the cells were present (Figure S4, Supplementary Information). A significant decrease in RNase activity (positive control) was observed in the presence of cells (Figure S4, Supplementary Information) also confirmed that A375 cells improved the ssRNA stability mainly via inhibiting the RNase activity of RNase-positive VTM. In addition, the nucleic acid content of A375 cells can act as a “carrier” to enhance the recovery and precipitation of ssRNA on magnetic beads, which can account for the lower *Ct* values observed for RNase-free VTM in the presence of cells.

### The effect of freeze-thaw cycles on SARS-CoV-2 qRT-PCR sensitivity

We studied the effect of multiple freeze-thaw cycles on the qRT-PCR detection sensitivity of ssRNA+cells sample (R+C) in HBSS and PBS-G VTM as the other VTM were observed to rapidly degrade the synthetic fragments as assessed after a single freeze-thaw cycle (Table 1). Similarly, the sensitivity of dsDNA plasmid (D) was studied in CDC and PBS-G VTM to test their cryopreservation properties. Further nucleic acid extractions were carried out after 3, 5, 7 and 10 freeze-thaw cycles for ssRNA+cells samples in HBSS and PBS-G and dsDNA samples in CDC and PBS-G given the above results. Each sample was extracted in triplicate and tested using qRT-PCR. When analysing qRT-PCR results, a cut-off point of cycle threshold (*Ct*) ≥ 35 was set for a negative sample.^34,35,28^

**Table 1.**
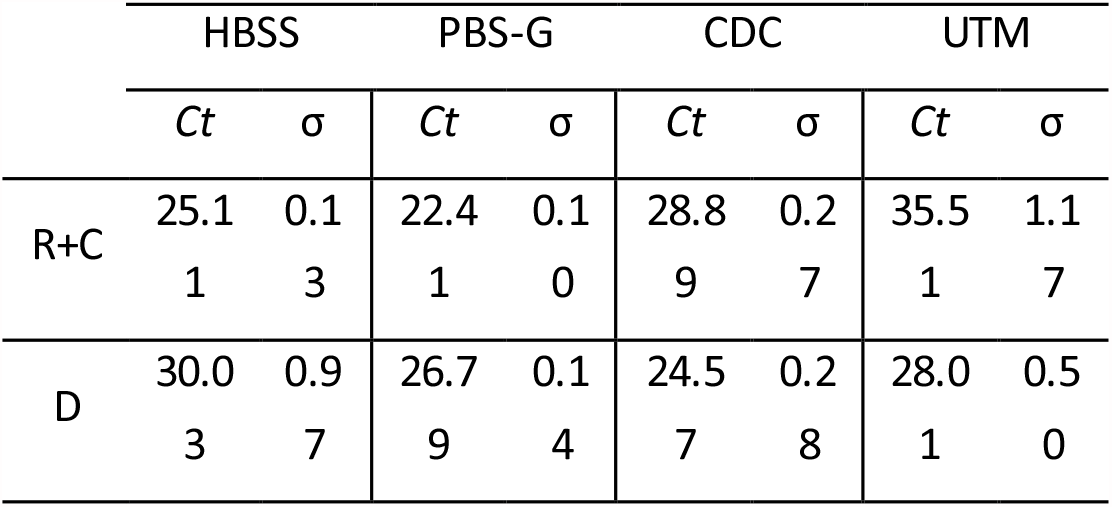
*Ct* values obtained in four different VTM after one freeze-thaw cycle for ssRNA+cells (R+C) and dsDNA (D) samples at 17,500 copies per qRT-PCR reaction.

The results for ssRNA+cells (R+C) and dsDNA (D) samples at 17,500 copies per qRT-PCR reaction extracted from the selected VTM are presented in Figure 3A and B, respectively. Two tailed t-tests (significance level of p < 0.01) were performed to compare the samples at each freeze-thaw cycle to their unthawed (“thaw 0”) counterpart. A statistically significant increase (p < 0.01) in *Ct* values after each freeze-thaw cycle was observed for ssRNA+cells in HBSS compared to their zero freeze-thaw counterpart (bars ‘H’ in Figure 3A). In contrast, freeze-thawing had no significant impact on the *Ct* values when PBS-G was used (bars ‘P’ in Figure 3A). For ssRNA+cells samples in HBSS, the *Ct* value increased by an average of 1.02 (σ = 1.23) (4.32%, σ = 5.50) for each subsequent freeze-thaw cycle condition tested. On the other hand, the average increase was 0.07 (σ = 0.19) (0.28%, σ = 0.75) per freeze-thaw cycle condition tested for the same sample type extracted from PBS-G, which suggests that the PBS-G could protect the ssRNA from degradation during freeze-thaw cycles better than HBSS. The findings suggest that the glycerol component of PBS-G VTM acts as a cryoprotectant and preserves the viral nucleic acid by preventing destructive ice crystals from forming during the freezing process.^36^ These crystals can damage the nucleic acid strands through a mechanical process known as shearing, in which one end of the strand is locked in an ice crystal whilst the other end is in the liquid phase that has not yet frozen, putting stress on the nucleic acid strand.^37^ Glycerol is thought to interfere with the solid structure of such crystals, preventing the formation of large ice crystals that may result in the physical stress and ultimately lead to shearing.^38^

**Figure 3.**
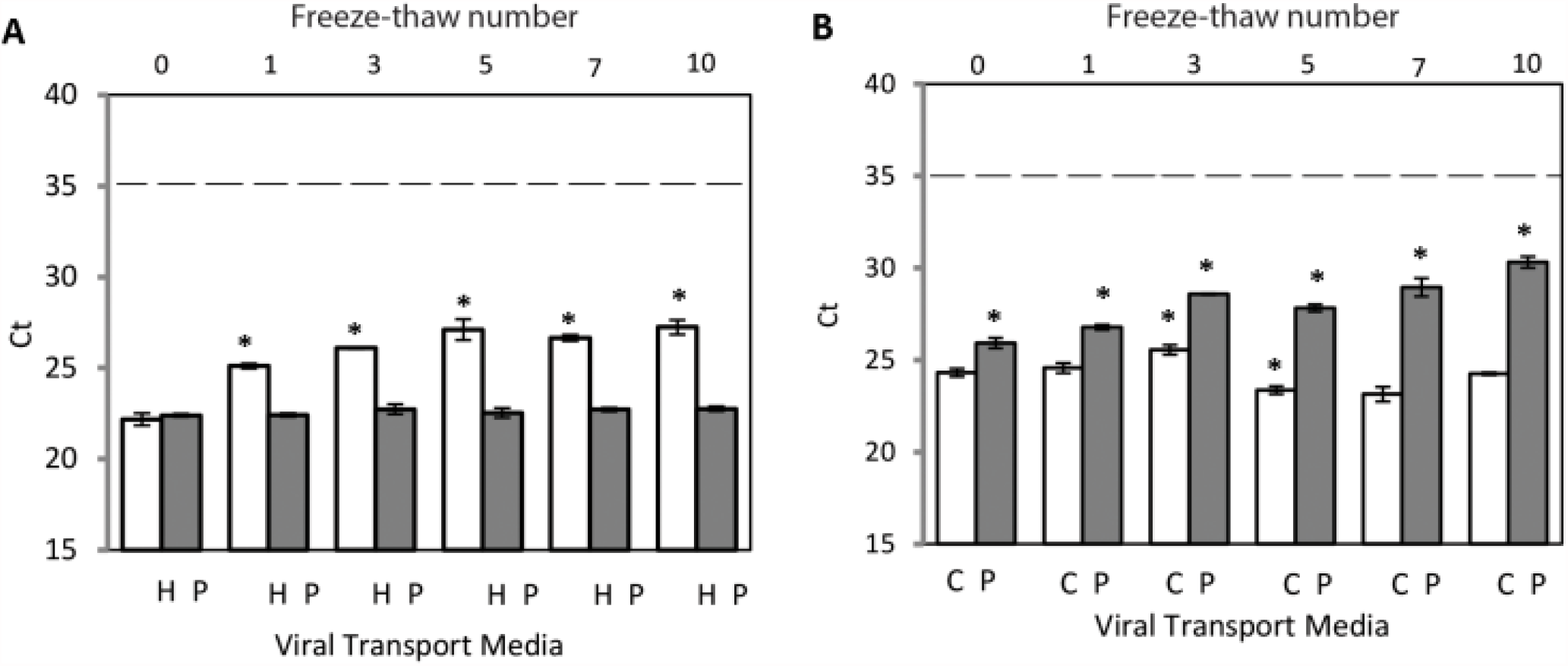
Impact of repeated freeze-thaw cycles on qRT-PCR measurements. **(A)** qRT-PCR results (*Ct*) plotted against ssRNA +Cell (R+C) samples extracted from HBSS (H) and PBS-G (P) VTM at 17,500 copies per qRT-PCR reaction after 0, 1, 3, 5, 7 and 10 freeze-thaw cycles. **(B)** qRT-PCR results (*Ct*) plotted against dsDNA (D) samples extracted from CDC (C) and PBS-G (P) VTM at 17,500 copies per qRT-PCR reaction after 0, 1, 3, 5, 7 and 10 freeze-thaw cycles.

Interestingly, the cryoprotective role of PBS-G was not evident for dsDNA samples, such that a significant increase in *Ct* values was observed (an average increase of 0.88 (σ = 0.97) (3.23%, σ = 3.51) for each subsequent freeze-thaw cycle condition tested) when the dsDNA plasmid underwent multiple freeze-thaw cycles in PBS-G (bars ‘P’ in Figure 3B). In CDC VTM, results changed significantly only after 3 (p= 0.0031) and 5 (p= 0.0057) freeze-thaw cycles (bars ‘C’ in Figure 3B), with an average decrease of 0.01 *Ct* (σ = 1.34) (0.07%, σ = 5.37) for each subsequent freeze-thaw cycle condition tested. Our observations suggest that the FBS component of CDC VTM protects the dsDNA plasmid during repeated freeze-thawing better than the glycerol component of PBS-G. Although not suitable for ssRNA samples due to RNase activity detected in its FBS component, we identified the CDC as a suitable VTM for the transport and storage of dsDNA.

Next, we studied the qRT-PCR detection sensitivity for varying viral copy numbers in repeated freeze-thaw experiments (Figure 4). We selected PBS-G and CDC VTM for ssRNA+cells and dsDNA plasmid samples, respectively. The tested viral copy numbers were 175, 1,750, and 17,500 per qRT-PCR reaction. As shown in Figure 4(a), the change in *Ct* values was not significant at different freeze-thaw cycles for all ssRNA+cells samples except for cycle 10 at 1,750 copies per qRT-PCR reaction. Similarly, *Ct* values for dsDNA plasmid in CDC VTM remained largely unaffected for all copy numbers except for freeze-thaw cycles 3 and 5 at 17,500 copies per qRT-PCR reaction (Figure 4B).

**Figure 4.**
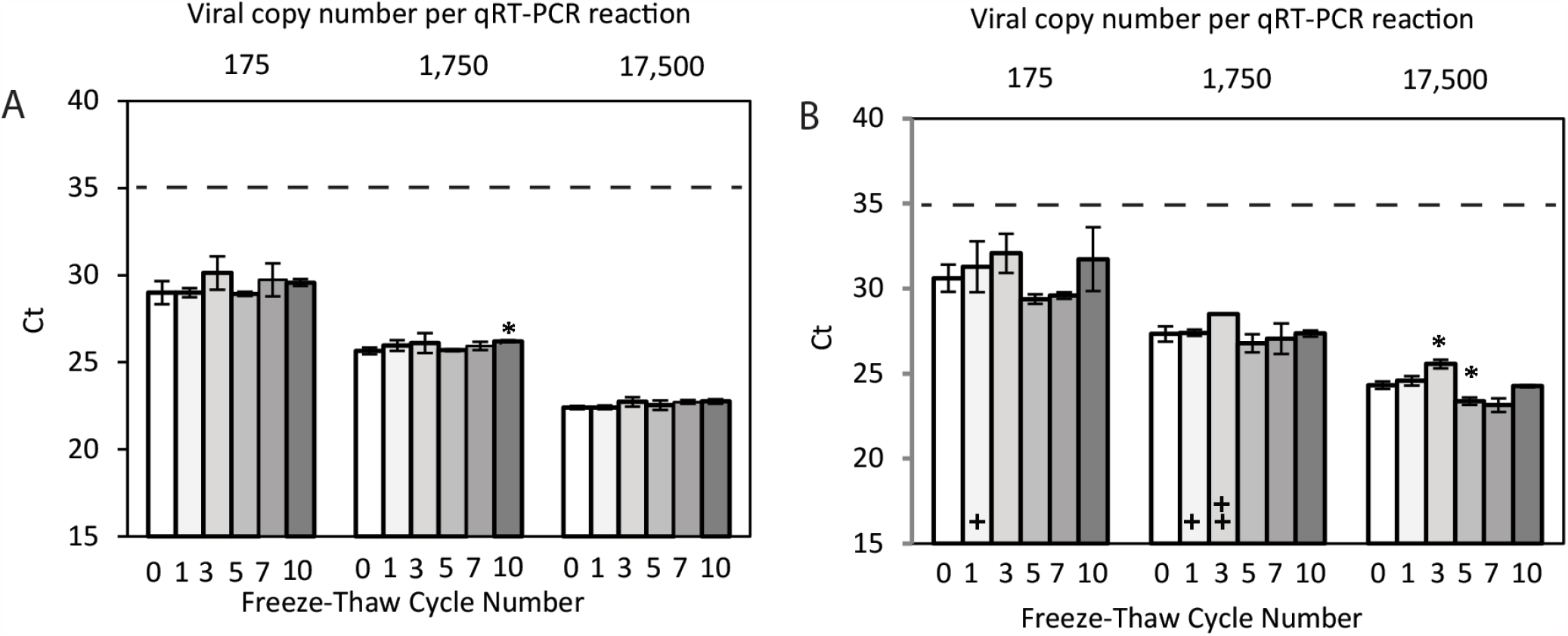
Impact of repeated freeze-thaw cycles on qRT-PCR measurements on synthetic SARS-CoV-2 controls over multiple concentrations. **(A)** ssRNA + Cell samples extracted from PBS-G VTM at 17,500, 1,750, 175 copies per qRT-PCR reaction after 0, 1, 3, 5, 7 and 10 freeze-thaw cycles. **(B)** dsDNA samples extracted from CDC VTM at 17,500, 1,750, 175 copies per qRT-PCR reaction after 0, 1, 3, 5, 7 and 10 freeze-thaw cycles. The number of triplicates per sample that failed to amplify is denoted by a cross (+) per failed triplicate in each sample bar i.e., one cross indicates one failed run and two crosses indicate two failed runs. Two tailed t-tests were used to compare each freeze-thaw cycle tested to their 0 freeze-thaw counterpart. Samples that produced a statistically significant difference (p < 0.01) are denoted by the presence of an asterisk (^*^).

The extraction of nucleic acids used in qRT-PCR in this study was based on the superparamagnetic beads that are larger than those in commercial extraction systems and have a higher speed of separation.^29,30^

The sensitivity of the qRT-PCR process based on this novel extraction system has been determined for clinical samples collected in PBS and benchmarked against commercial platforms. ^39^ The diagnostic sensitivity of this novel system exceeded several commercials platforms by 1-2 *Cts* resulting in lower level of false negatives at low viral copy numbers.

Our data are in line with results obtained from studies on freeze-thaw cycles of COVID-19 samples. Dzung et al.^27^carried out extractions on patient-derived samples after up to 15 freeze-thaw cycles and implemented qRT-PCR analysis. It was found that although the *Ct* increase from 1 to 5 and from 5 to 10 freeze-thaw cycles was statistically significant, the change was not high enough to prevent the viral detection as the virus was still detectable after 15 freeze-thaw cycles. The reported average *Ct* increase was 0.106 to 0.197 per freeze-thaw cycle, which are consistent with our findings. Notably, the VTM used in this study was created using a modified version of the VTM protocol from Institute of Medical Virology (University of Zurich, Zurich, Switzerland) and contained fetal calf serum. However, in our study, the freeze thaw cycles had less of an influence on the *Ct* values for ssRNA in PBS-G and dsDNA in CDC. This is to be expected as we specifically selected the VTM that we thought would withstand the freeze-thaw cycles best with the materials we used, and one of our materials was a dsDNA plasmid which is inherently more stable. As well as this, due to the use of synthetic nucleic acids, we were able to select the concentration of viral target to test. So, whilst Dzung et al. noted that they only tested samples with a moderately high viral load and therefore could not investigate the effects the number of freeze-thaw cycles would have on a samples with lower viral loads^27^, we were able to show that even at a lower concentration of 175 viral copies per qRT-PCR reaction there was no significant difference in *Ct* between samples that had not been freeze-thawed compared to those that underwent 10 freeze-thaw cycles.

## Conclusions

The COVID-19 pandemic has presented a worldwide challenge to healthcare systems and particularly to analytical laboratories that have had to execute hundreds of millions of new molecular diagnostics tests. The results presented in this article highlight the important role that VTM selection has on the sensitivity of qRT-PCR detection of RNA viruses. It is apparent that simple buffers are preferable to more complex ones that include serum, which have been shown to be contaminated with RNases. Figure 5 illustrates the mechanism that we believe leads to the degradation of qRT-PCR signal from VTM containing RNases.

**Figure 5.**
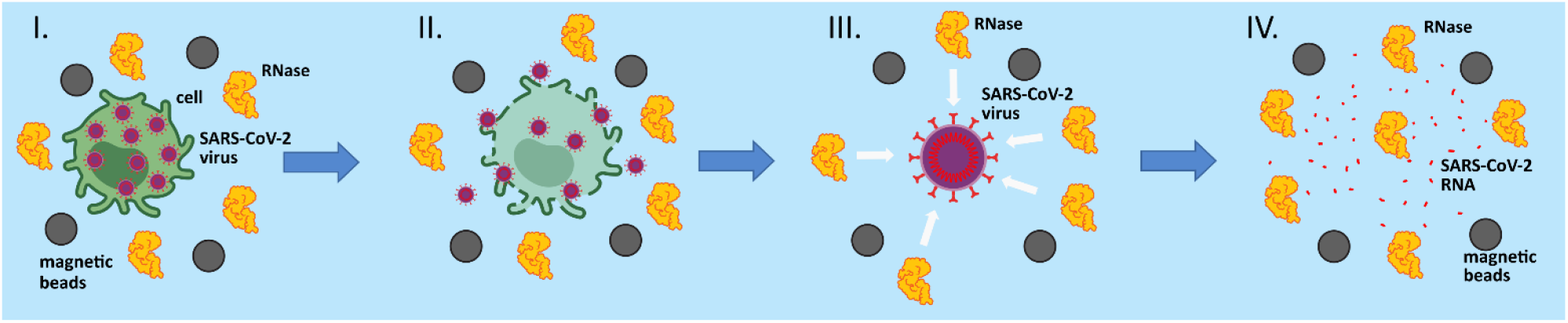
A graphical illustration of RNA extraction in the presence of a viral transport medium that has been contaminated with RNase activity. (**I-III**) Lysis of cells and viral particles in the presence of denaturants and detergents. Residual RNase activity from the VTM remains despite the presence of denaturants and DTT. (**IV**) SARS-CoV-2 RNA that is released from viral particles is degraded by RNases rendering it unsuitable for capture, purification and detection by molecular diagnostic assays.

This article also demonstrated that multiple freeze-thaw cycles do decrease the sensitivity of qRT-PCR detection of RNA viruses in some VTM. Specifically, little change in sensitivity was observed for samples collected in RNase-free VTM and those containing inexpensive cryopreservatives, such as glycerol. This is a particularly important observation for COVID-19 testing in resource-limited environments, where the cold chain and logistics can be difficult to maintain and are more easily overextended.

An unexpected observation made in this study was that cellular material from cultured A375 cells enhanced the sensitivity of qRT-PCR detection of RNA viruses. It appears there is one or more compounds in the nuclear or cytosolic extracts of these epithelial cells that stabilize the viral RNA and thus lead to the enhanced performance of the molecular diagnostics assays.

The results presented in this article also suggest that DNA viruses are likely to be less sensitive to the composition of VTM than RNA viruses. It appears that simple buffers with cryopreservatives allow efficient storage and transfer of samples with minimal loss of viral copy numbers. However, sources of DNases need to be carefully identified and monitored in these VTM as well.

## Author Contributions

**CH, SH, and NF** Methodology, validation, analysis, investigation, visualization, and writing. **PL, JO and CM** Resources. **OT, MC and GL** Concept, writing, supervision

## Acknowledgements

This work was supported by the Science Foundation of Ireland (15/IA/3127 and 20COV0054) and the EU Cordial-S programme under grant agreement No 101016038. We would like to acknowledge Connor Field, Tofie Ajomale, Maria Materia, Ben Lindsay, Glenna Swindon, Sophie O’Riley, Virginie Gautier and Paddy Mallon.

## References

1. World Health Organisation (WHO), 2021. COVID-19 Weekly Epidemiological Update. Emergency Situational Updates. [online] Available at: <https://www.who.int/emergencies/diseases/novel-coronavirus-2019/situation-reports> [Accessed 17 May 2021].

2. T. Ji, Z. Liu, G. Wang, X. Guo, S. Akbar Khan, C. Lai, H. Chen, S. Huang, S. Xia, B. Chen, H. Jia, Y. Chen and Q. Zhou, Biosensors and Bioelectronics, 2020, 166, 112455.

3. Centers for Disease Control and Prevention, 2020. CDC 2019-Novel Coronavirus (2019-nCoV) Real-Time RT-PCR Diagnostic Panel. Revision 6. [PDF] Available at: <https://www.fda.gov/media/134922/download> [Accessed 30 April 2021].

4. Y. Pan, L. Long, D. Zhang, T. Yuan, S. Cui, P. Yang, Q. Wang and S. Ren, Clinical Chemistry, 2020, 66, 794–801.

5. Y. Hsin-Chih, E. Simone, C. Zhang and T.H. Wang S, 17th IEEE International Conference on Micro Electro Mechanical Systems. Maastricht MEMS 2004 Technical Digest, 2004, 371–374, doi: 10.1109/MEMS.2004.1290599.

6. E. Surkova, V. Nikolayevskyy and F. Drobniewski, The Lancet Respiratory Medicine, 2020, 8, 1167–1168.

7. S. Klein, T.G. Müller, D. Khalid, V. Sonntag-Buck, A.M. Heuser, B. Glass, and P. Chlanda, MedRxiv. 2020, https://doi.org/10.1101/2020.07.08.20147561

8. K.A. Melzak, C.S. Sherwood, R.F.B., Turner, and C.A. Haynes, Journal of Colloid and Interface Science, 1996, 1996, 635–644.

9. P.E. Vandeventer, J.S. Lin, T.J. Zwang, A. Nadim, M.S. Johal, and A. Niemz, Journal of Physical Chemistry B, 2012, 116, 5661–5670.

10. D. Relova, L. Rios, A. Acevedo, L. Coronado, C. Perera and L. Pérez, Veterinary Sciences, 2018, 5, 19.

11. Y. Zhang, C. Wang, M. Han, J. Ye, Y. Gao, Z. Liu, T. He, T. Li, M. Xu, L. Zhou, G. Zou, M., Lu, and Z. Zhang, Virologica Sinica, 2020, 35, 758–767.

12. T. Kwon, N.N. Gaudreault, and J.A. Richt, Pathogens, 2021, 10, 540. https://doi.org/10.3390/pathogens10050540

13. Y. Nishibata, S. Koshimoto, K. Ogaki, E. Ishikawa, K. Wada, M. Yoshinari, Y. Tamura, R. Uozumi, S. Masuda, U. Tomaru and A. Ishizu, Pathology, research and practice, 2021. https://doi.org/10.1016/j.prp.2021.153381

14. W. Wang, Y. Xu, R. Gao, R. Lu, K. Han, G Wu, and W. Tan, JAMA, 2020, 2020, 1843–1844.

15. F. Gao, L. Tao, X. Ma, D. Lewandowski and Z. Shu, Biopreservation and Biobanking, 2020, 18, 511–516.

16. R.M. Martinez, Clinical Microbiology Newsletter, 2020, 42, 121–127.

17. Centers for Disease Control and Prevention, 2020. Preparation of Viral Transport Medium. Standard Operating Procedure SOP#: DSR-052-02. https://www.cdc.gov/coronavirus/2019-ncov/downloads/Viral-Transport-Medium.pdf

18. World Health Organisation, ‘Collecting, preserving and shipping specimens for the diagnosis of avian influenza A(H5N1) virus infection Guide for field operations’. 2006, Annex 8, pp. 42–43.

19. A.A. Rogers R.E. Baumann G.A. Borillo R.M. Kagan H.J. Batterman, M.M. Galdzicka, and E.M. Marlowe J Clinical Microbiology, 2020, 58, 708–720.

20. S. Summer, R. Schmidt, A. Herdina, I. Krickl, J. Madner, G. Greiner, F. Mayer, N. Perkmann-Nagele and R. Strassl R., MedRxIV, 2020, doi: https://doi.org/10.1101/2020.07.21.20158154

21. P.D. Kirkland and M.J. Frost Pathology 2020, 52, 811–814.

22. A. Chin, J. Chu, M. Perera, K. Hui, H. Yen, M. Chan, M. Peiris, and L. Poon, The Lancet Microbe, 2020, 1, p.e10.

23. H. Aboubakr, T. Sharafeldin and S. Goyal, Transboundary and Emerging Diseases, 2020, 68, 296–312.

24. W. Shao, S. Khin and W. Kopp, Biopreservation and Biobanking, 2012, 10, 4–11.

25. K. Yu, J., Xing, J. Zhang, R. Zhao, Y., Zhang and L. Zhao, Cell and Tissue Banking, 2017, 18, 433–440.

26. D. vanBockel, C. Munier, S. Turville, S.G. Badman, G. Walker, A.O. Stella, A. Aggarwal, M. Yeang, A. Condylios, A. Kelleher, T.L., Applegate, A. Vallely, D. Whiley, W. Rawlinson, P. Cunningham, J. Kaldor and R. Guy, Viruses, 2020, 12, 1208.

27. A. Dzung, P. Cheng, C. Stoffel, A. Tastanova, P. Turko, M. P. Levesque, P.P. Bosshard, The Journal of Molecular Diagnostics, 2021, 220, 153381.

28. J.J.J.M. Stohr, M. Wennekes, M. vanderEnt, B.M.W. Diederen, M.F.Q. Kluytmans-van den Bergha, A.M.C. Bergmans, J.A.J.W. Kluytmans, and Suzan D. Pas, J. Clin. Virol., 2020, 133, 104686.

29. H. Shang, W.S. Chang, S. Kan, S.A. Majetich, G.U. Lee, Langmuir, 2006, 22, 2516–2522.

30. J.J. O’Mahony, M. Platt, D. Kilinc, and G. Lee, Langmuir, 2013, 29, 2546–2553.

31. Life technologies, 2013, RnaseAlert QC System v2: User Guide.

32. R. Wölfel, V.M. Corman, W. Guggemos, M. Seilmaier, S. Zange, M.A. Müller, D. Niemeyer, T.C. Jones, P. Vollmar, C. Rothe, M. Hoelscher, T. Bleicker, S. Brünink, J. Schneider, R. Ehmann, K. Zwirg lmaier, C. Drosten, and C. Wendtner, Nature, 2020, 581, 465–469.

33. A. Granados, A. Petrich, A. McGeer and J.B., Gubbay, Journal of Virology Methods, 2017, 247, 45–50.

34. M. Tom and M. Mina, Clinical Infectious Diseases, 2020. 71, 2252-2254.

35. K.M. Mokhtar, medRxiv 2020 doi: 10.1101/2020.11.20.20235390

36. F. Johnson, Clinical Microbiology Reviews, 1990, 3, 120–131.

37. J. Brunstein, 2015. Freeze-thaw cycles and nucleic acid stability: what’s safe for your samples?

38. B. Röder, K. Frühwirth, C. Vogl, M. Wagner, P. Rossmanith, Journal of Clinical Microbiology, 2010, 48, 4260–4262

39. S. O’Reilly, N. Feely, P. Li, R. Murtagh, J. Urquiza, A.G. Leon, L.E. McLoughlin, G. Swinand, C. Holohan, V. D’Autume, S. McDermott, P. Mallon, G.U Lee, and V. Gautier, COVID-19 RAPID RESPONSE: Development of a high-throughput viral RNA extraction platform to sustain COVID-19 molecular diagnostics, presented as part of the SFI Summit, Dublin, Ireland, October 2020.

