## Supplementary Information for "Influence of viral transport media and freeze-thaw cycling on the sensitivity of qRT-PCR detection of SARS-CoV-2 nucleic acids"

**RNase Activity Assay**

The RNaseAlert QC system V2 Kit from Thermo Fisher was used to test the solutions for RNase activity using the supplied protocol. Samples, positive and negative controls were tested in duplicates. RNase-free, filtered pipette tips were used throughout. Recommended fluorometer settings from the RNaseAlert QC system V2 protocol were followed. The temperature, focal height and gain settings were 37 ^o^C, 3.4, and 50%, respectively. The excitation/emission wavelengths were 490/520nm. The well scan setting was set to orbital and a mixing step was selected for 5 seconds before plate reading at 500rpm. The time interval for data collection was set to 1.5 minute intervals for 20-30 cycles i.e. 30- 45 minutes.

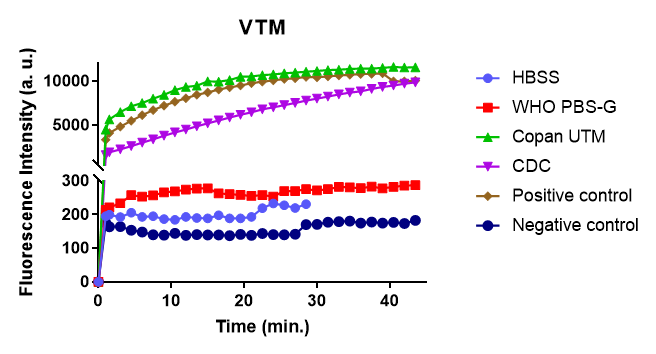

**Figure S1.** RNase activity assay (RNaseAlert QC system V2) performed on different VTM used in this study in comparison to positive and negative controls. UTM (green) and CDC (purple) displayed high levels of RNase activity that was comparable to positive control (brown). HBSS (blue) and PBS-G (red) presented as negative for RNase activity, comparable to negative control (navy). Positive control: RNase A, Negative control: nuclease-free water (black).

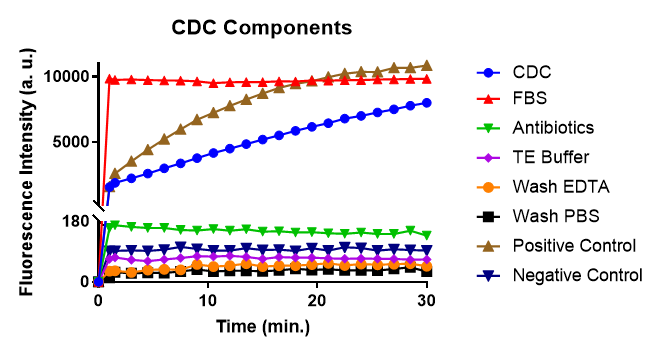

**Figure S2.** RNase activity assay (RNaseAlert QC system V2) performed on the different components of CDC (blue) that displayed a high RNase activity. All CDC components were RNase-free except for FBS (red), which displayed a higher RNase activity compared to the positive control (brown). Positive control: RNase A, Negative control: nuclease-free water (navy).

**Figure S3.** RNase activity assay (RNaseAlert QC system V2) performed on the various components of V-REK nucleic acid extraction kit used in this study in comparison to positive (brown) and negative (navy) controls. All components tested negative for RNase activity. Positive control: RNase A, Negative control: nuclease-free water.
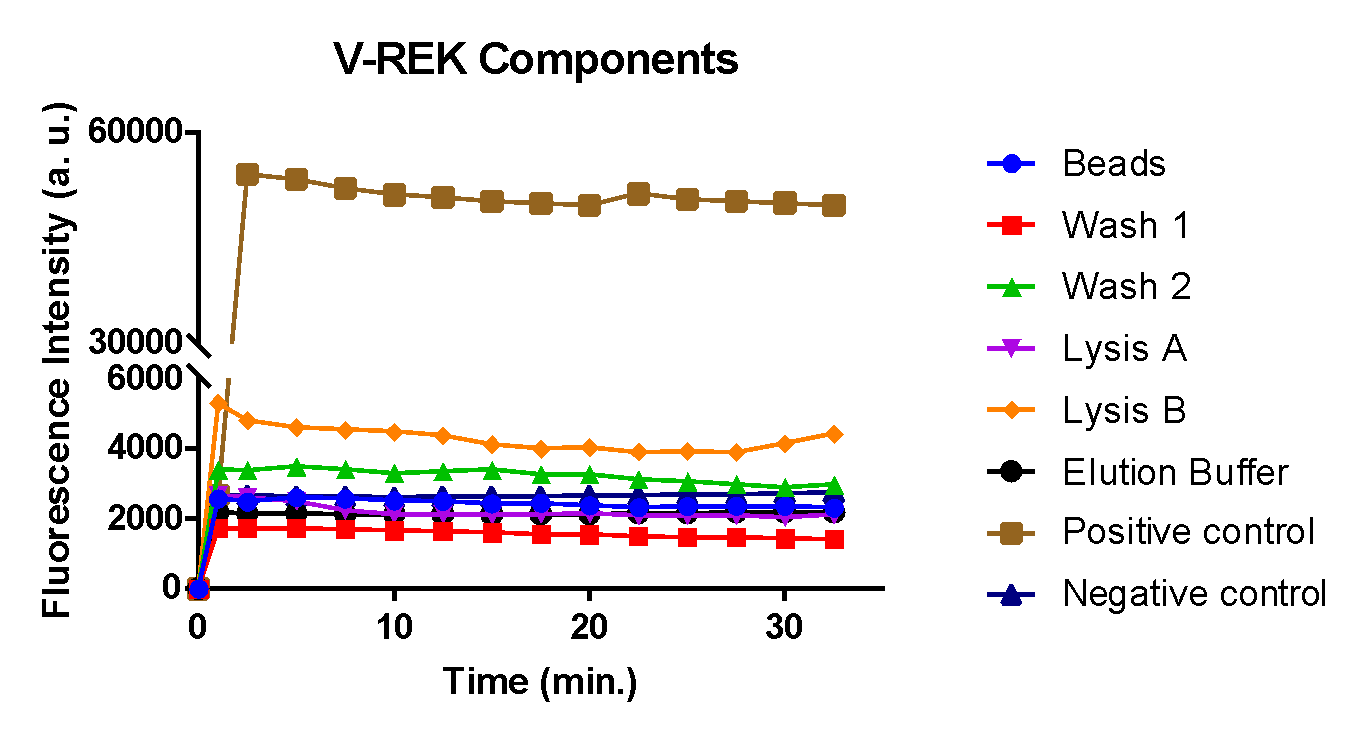

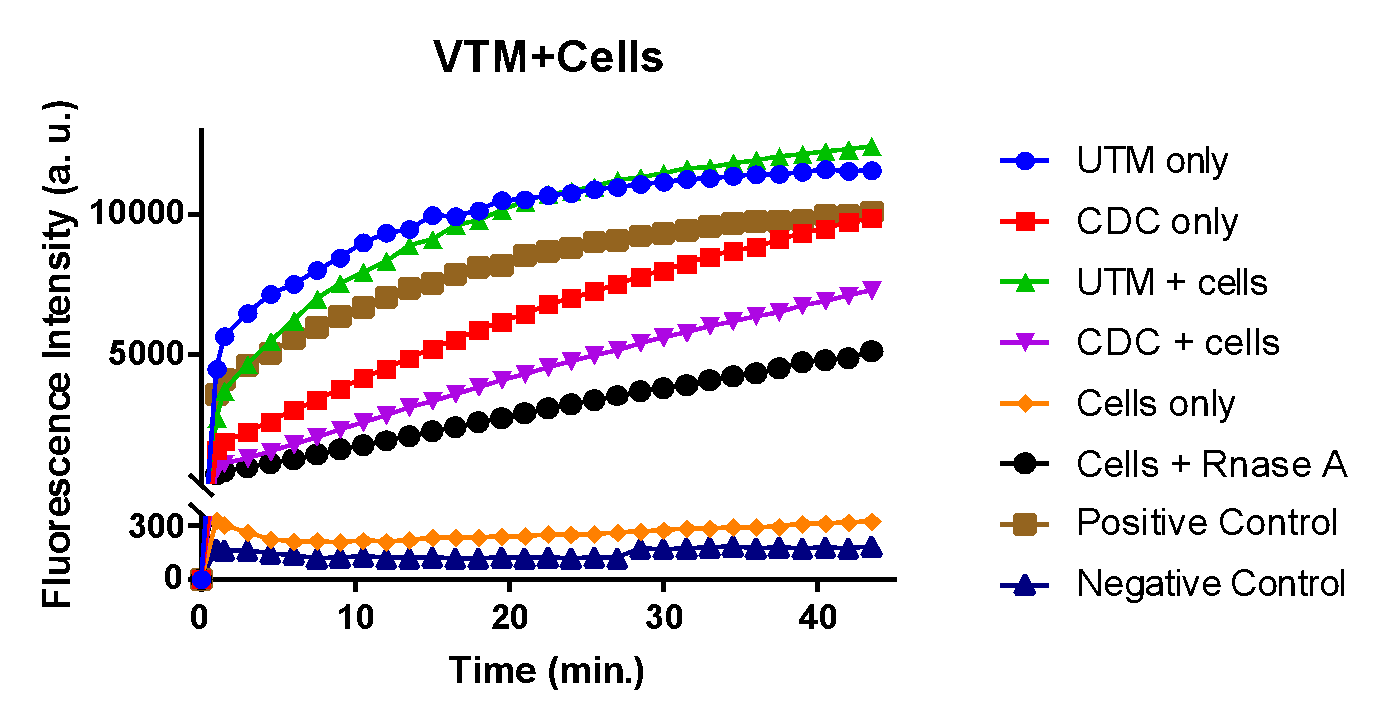

**Figure S4.** RNase activity assay (RNaseAlert QC system V2) performed on the RNase-positive VTMs in the presence of A375 cells. The RNase activity in UTM (blue) and CDC (red) decreased in the presence of cells (UTM+cells, green; CDC+cells, purple). The positive control RNase A (brown) also displayed a reduced activity when tested in the presence of A375 cells (Cells+Rnase A, black). RNase activity of A375 cells (Cells only, orange) was comparable to the negative control (nuclease-free water, navy).
